# Mobius Assembly for Plant Systems highlights promoter-coding sequences-terminator interaction in gene regulation

**DOI:** 10.1101/2024.07.10.602858

**Authors:** Elif Gediz Kocaoglan, Andreas Andreou, Jessica Nirkko, Marisol Ochoa-Villarreal, Gary Loake, Naomi Nakayama

**Author notes:** These authors contributed equally (Elif Gediz Kocaoglan, Andreas Andreou). Corresponding author: Naomi Nakayama –.

## Abstract

Plants are the primary biological platforms for producing food, energy, and materials in agriculture; however, they remain a minor player in the recent synthetic biology-driven transformation in bioproduction. Molecular tools and technologies for complex, multigene engineering in plants are as yet limited, with the challenge to enhance their stability and predictivity. Here, we present a new standardized and streamlined toolkit for plant synthetic biology, Mobius Assembly for Plant Systems (MAPS). It is based on small plant binary vectors pMAPs, which contain a fusion origin of replication that enhances plasmid yield in both *Escherichia coli* and *Rhizobium radiobacter*. MAPS includes a new library of promoters and terminators with different activity levels; part sizes were minimized to improve construct stability and transformation efficiency. These promoters and terminators were characterized using a high-throughput protoplast expression assay. We observed a significant influence of terminators on gene expression, as the strength of a single promoter can change more than seven-folds in combination with different terminators. Changing the coding sequence changed the relative strength of promoter and terminator pairs, thus uncovering combinatorial gene regulation among all parts of a transcriptional unit. We further gained insights into the mechanisms of such interactions by analyzing RNA folding, with which we suggest a design principle for more predictive and context-independent genetic parts in synthetic biology of plant systems and beyond.

## INTRODUCTION

Throughout the history of civilization, plants have been the primary culturing platform for food, energy, materials, and drugs. However, in synthetic and engineering biology applications, plants have remained a minor vehicle for bioproduction with untapped potential. Consequently, the molecular tools and technologies for rational engineering of plant systems are still limited compared to their microbial and mammalian counterparts. With the increasing world population and ongoing climate crisis, considerately designed plant engineering holds the key to a sustainable future, creating transformative solutions for a wide range of applications from agriculture to environmental regeneration^1–4^. From enhancing crop yields, quality, and resilience to improving carbon fixation and biofuel efficiency, synthetic biology-based plant engineering is becoming a powerful catalyst to futureproof our society and planet^5–7^.

Synthetic biology enables complex pathway engineering necessary for modifications of biological metabolisms, cellular and developmental pathways, or responses to pathogens and diseases. The foundation of synthetic biological genetic engineering is the engineering principles, such as standardization, modularity, and design-build-test cycles^8^. However, as the field matures past its first two decades, biology has often defied these fundamental assumptions. The concept of orthogonality – context-independent, universal functions of genetic parts – is routinely challenged, uncovering new insights and understanding of how biological systems interact^9^. As this can be the case for just a single gene, it poses a significant hurdle when working with the many genes necessary for pathway and network engineering.

The development of synthetic biology tools and technologies for multigene engineering starts with creating a collection (library) of standardized genetic parts. Standardized parts (e.g., Phytobricks)^10^ include promoters, coding sequences, terminators, and functional protein tags, which together form transcriptional units (TUs). Promoters, which contain the transcription start site (TSS) where RNA polymerase binds, are well established for their instructive roles in regulating gene activity. In comparison, terminators, typically found at the end of a gene, are often underestimated in their regulatory roles. However, they have been shown to influence transcription by controlling transcription arrest, mRNA stability, protecting genes against silencing and tuning other transcription functions^11^.

The most commonly used promoters and terminators in plant genetic engineering originally stemmed from Cauliflower Mosaic Virus (*35S*), *Rhizobium* opine genes (*NOS, MAS,* and *OCS*), and more recently, plant genomes (e.g., *UBQ10* promoter and *HSP* terminator from *Arabidopsis thaliana* or hereafter Arabidopsis). The shortage of well-characterized regulatory parts results in repeated use of the same sequences in multigene construction, increasing the possibility of homologous sequence-dependent recombination and gene silencing^12,13^. In addition, there is a need for shorter standard parts, especially when multiple genes are delivered in a single vector, to reduce the size of the construct. Large constructs tend to make plasmids structurally unstable^14^ and decrease the efficiency of *Rhizobium*-mediated transformation^15^. Therefore, creating novel and shorter promoter and terminator parts will enrich the plant genetic engineering toolkit available to the community and increase our capacity to create genetic modifications.

Once standard genetic parts are made, their activity level should be assessed. This typically involves combining a promoter and terminator pair with a reporter protein coding sequence and quantifying the reporter expression as output. While promoters are generally more well characterized, recently, attention has also been directed towards terminators; diverse arrays of terminators have been characterized in bacteria^16^ and yeast^17^. For plant systems, the number of available terminator parts was limited for a long time, with their characterization typically performed using the *Nicotiana benthamiana* leaf infiltration system^18,19^. It is worth noting that different species, or even different expression systems within the same species, may exhibit varying levels of expression for the same TU^18,19^. Therefore, to achieve higher stability and predictability, new standard parts should be characterized in different combinations and chassis environments to account for potential context dependency.

As the number of constructs to build and characterize increases, molecular construction needs to become more efficient. Golden Gate is a widely used technology for DNA assembly that utilizes Type IIS restriction endonucleases that cut DNA outside their recognition sequences, allowing the introduction of short ‘sticky end’ sequences (syntaxis) that help orient and combine multiple DNA fragments in an intended order. Many Golden Gate frameworks have been developed for diverse species^20–23^; recent developments in the plant field include Loop Assembly^24^, an extension of the MoClo toolkit for plants^25^, and Joint Modular Cloning^18^. We have previously developed a new Golden Gate framework^26^, named Mobius Assembly, for its iterative cloning strategy between two vector sets; it allows theoretically infinite assembly of up to four DNA fragments each time. It is designed to be simple, versatile, and efficient (e.g., less need for domestication), complying with the universal and popularly employed the syntaxis design Phytobrick^27^.

In plant transgenic engineering, binary vectors house transgenes and facilitate their delivery and incorporation into the recipient genomes. Although functional, these vectors could be improved to enhance transformation efficiency and stability. A typical *Rhizobium*-mediated transformation vector includes T-DNA borders^28^, replication functions for *E. coli* and *Rhizobium*, and selection markers. Plasmid instability is a significant challenge, especially for plant systems, as vectors must be stable in both *E. coli* and *Rhizobium*. Factors such as size^14^, copy number^29^, direct repeats12 and inverted repeats^30^ affect plasmid stability. Traditional vectors are bulky (>6 kb), but smaller backbones like pGreen (2.5 kb^31^), pLSU (4.6 kb^32^), and pLX (3.3 kb^33^) have been developed to reduce instability. The pLX vectors use a modular design with minimal functional parts and stability features, improving binary vector construction.

Here, we report the adaptation of Mobius Assembly for plant species engineering: Mobius Assembly for Plant Systems (MAPS). The MAPS toolkit includes new, compact plant binary vectors (pMAPs). These vectors are 3.6 kb in size, structurally stable through molecular construction, and suitable for transformation methods requiring high plasmid DNA amounts. MAPS also features bioluminescence and fluorescent protein reporters, along with new standardized promoter and terminator parts for a range of gene expression strengths. In high-throughput transient expression tests with *Arabidopsis thaliana* protoplasts, MAPS promoters effectively drove gene activation gradients. We also demonstrated that terminators can be used to control gene expression levels. The strength of the promoter and terminator parts depended on their combinations, and such interactions extended to the coding sequence selection. These findings facilitate a bottom-up understanding of gene regulation and the design of context-independent molecular constructs with more predictable outputs.

## RESULTS

### Development of Mobius Assembly for Plant Systems (MAPS)

MAPS is an extension of our molecular cloning framework Mobius Assembly^26^ to enable transformation and expression of transgenes in plant systems. MAPS has a universal acceptor vector (mUAV) at Level 0 to house a standard part in the Phytobrick format. Level 0 parts are combined at Level 1, and up to four Level 1 constructs are then combined to make Level 2 constructs. MAPS follows a linear cloning strategy until Level 2 and then iterates between two cloning levels (Level 1 and Level 2) for quadruple augmentation of cloning units each time (**Fig. 1a**). Using the rare cutter *AarI (PaqCI),* as opposed to frequently used restriction enzymes that recognize shorter sequences (e.g., *BsmBI* or *BpiI*), reduces the need for removing internal restriction sites (*i.e.,* domestication).

**Figure 1.**
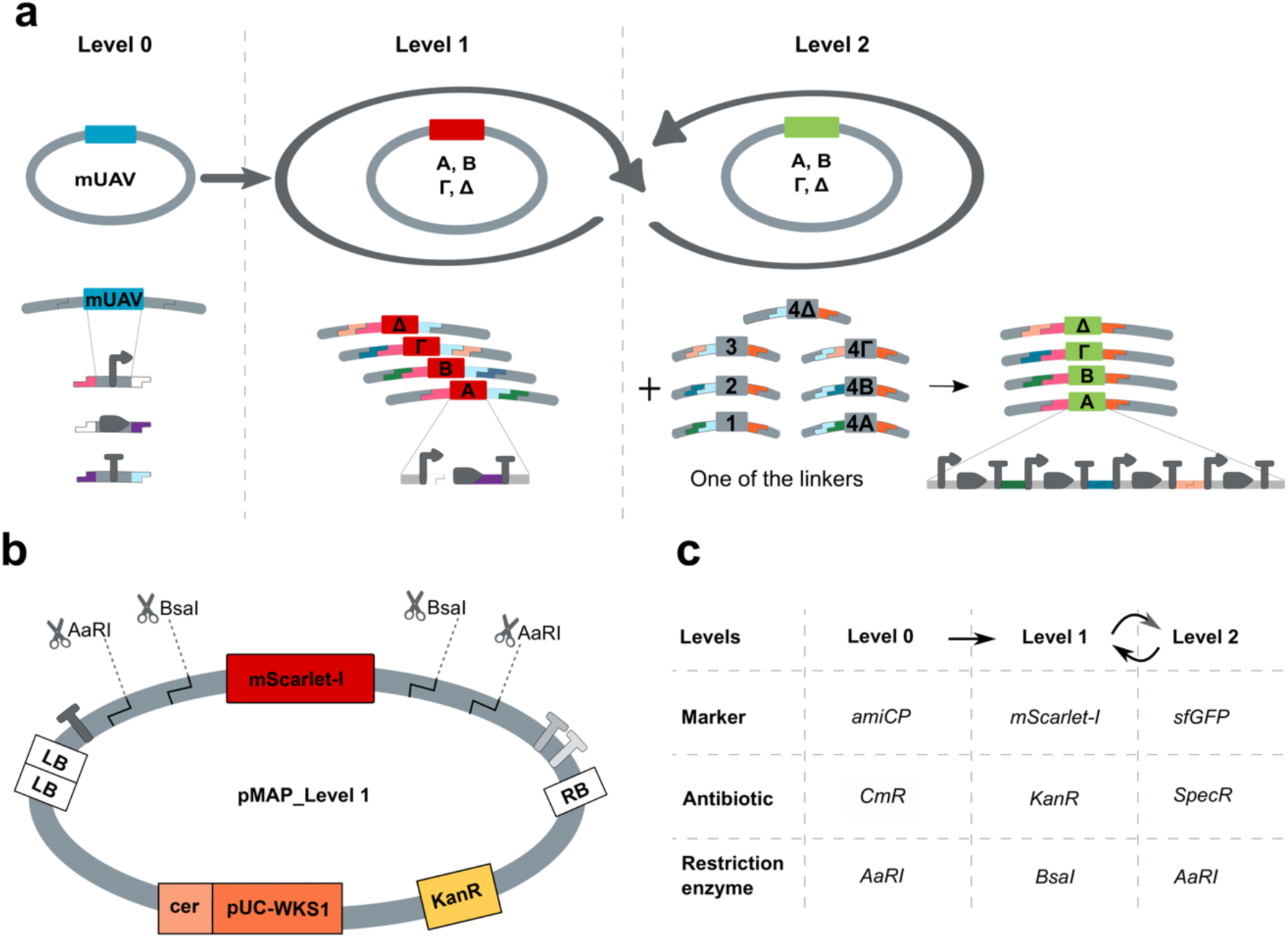
Mobius Assembly for Plant Systems (MAPS) vector structure and workflow. **a** Schematics of how Mobius Assembly operates. MAPS is an adaptation of the Mobius Assembly for plant systems based on small binary vectors intended for plant transformation. The core vector toolkit is comprised of one part storage vector in Level 0 (mUAV), four Level 1 Acceptor Vectors, fourLevel 2 Acceptor Vectors, and seven Auxiliary plasmids providing linkers. The vectors release the insert with *BsaI* in Level 1 cloning and the rare cutter *AarI (PaqCI)* in Level 0 and Level 2 cloning. **b** Plasmid design of MAPS vectors, showing a Level 1 plasmid as an example. At Level 2, *BsaI* and *AaRI* (*PaqCI*) cut sites swap places. **c.** Summary of the markers, antibiotic resistance and RE enzymes used for Mobius Assembly at different Levels. *CmR* is a gene conferring the resistance for chloramphenicol, *KanR* for kanamycin and *SpecR* for spectinomycin. For simplicity, the overhangs carried from Level 0 are not shown in the Level 2 construct.

Initially, we developed pGreen-based vectors but encountered issues with large constructs consistent with reported instability issues^34,35^. To address this, we created a new small plant binary vector called pMAP, based on the pLX architecture (**Fig. 1b**). This vector is suitable for transient expression in protoplasts, cell culture, tissues/organs, and whole-plant stable transformation. We devised a new origin of replication by fusing pWKS1 and pUC19 Ori. The pWKS1 Ori, derived from *Paracoccus pantotrophus* DSM 11072^36^, is functional in *Rhizobium* but not in *E. coli*, so we fused it with the minimal stable pUC Ori from pUC19. pMAPS also has two Left Border (LB) sequences instead of one. *Rhizobium* gene transfer is from the Right Border (RB) to the LB, and having two LBs suppresses backbone transfer to the plant genome^37^. Another feature of pMAPS is that terminators flank LB and RB sequences to isolate transgene activity from the plasmid backbone. Colourific markers, antibiotics selections, and restriction enzymes used for Mobius Assembly at different levels are summarized in **Fig. 1c**.

MAPS vector toolkit consists of a core set of pMAP cloning/destination vectors (Level 1 and Level 2 Acceptor Vectors, four variations Α-Δ for each level), which have a fusion origin of replication to replicate in *E. coli* and *Rhizobium*. The mUAV and the seven Auxiliary plasmids are also included, as described in the original Mobius Assembly kit^26^ . The MAPS toolkit also contains a selection of plant promoters, terminators, antibiotic resistance genes, and visible reporter genes (bioluminescence and fluorescent proteins). All MAPS plasmids are listed in **Supplementary Fig. S1; Supplemental Tables S1, S2** and available through AddGene (https://www.addgene.org/browse/article/28211394/).

Reflecting on user feedback, two specific improvements were made to improve the Mobius Assembly vectors. A few users of the original Mobius Assembly kit indicated that the chromoprotein selection had been lost in their clones. Upon investigation, we observed independent events of transposon insertion in the promoter of the chromoprotein genes, which we hypothesized is a response to the stress imposed by chromoprotein production (**Supplementary Fig. S2).** To solve the problem, we replaced the marker chromogenic protein (spisPINK) with a red fluorescent protein (mScarlet-I). (**Supplementary Fig. S2).** We also noticed some Mobius Assembly clones showed growth retardation in the selection media and identified the instability was caused by plasmid dimerization. Therefore, we introduced a 240-bp *cer* domain, which recognizes dimers and triggers recombination help keep the plasmids in the monomeric state (**Supplementary Fig. S3**)^38^.

Combinatorial DNA libraries are crucial for part characterization, as well as in applications such as biosynthetic pathway optimization^39^, but their manual construction is time-consuming and resource-intensive. To aid combinatorial library construction, we developed the ‘MethylAble’ feature in Mobius Assembly, allowing standard part variants (Level 0) to be introduced to specific sites in single or multi-gene constructs (**Supplementary Fig. S4)**. MethylAble utilizes the DNA methylation sensitivity of *BsaI* to mask its recognition sites by cytosine methylation during Level 1 cloning. We designed an *amilCP* expression cassette with divergent and convergent *BsaI* recognition sites, where CpG methylation blocks *BsaI* digestion only at divergent sites, allowing insertion of Level 0 parts into premade Level 1 constructs. Correct constructs show a purple color from *amilCP* until the Level 0 parts replace the cassette. As a proof of concept, the MethylAble protocol was used to build the library of the three inducible promoters (see below), each of which was combined with the 14 terminator coparts, making 42 constructs in total (**Supplemental Table S3**). MethylAble presents a novel strategy to create construct libraries and can be implemented in all Golden Gate frameworks in which *BsaI* enzyme is used, not only in Mobius Assembly.

### Designing, building, and testing MAPS promoter/terminator standard parts

To select new ‘constitutive’ promoter and terminator parts, we chose ubiquitously expressed genes that are likely to have strong expression in different tissue types **(Supplementary Table S2).** The promoters and terminators were characterized with a transient gene expression assay based on Arabidopsis mesophyll protoplasts and PEG transformation. We optimized the parameters throughout the protocol based on^40^ to improve the transformation efficiency and reproducibility. We were able to reach up to 70% transformation efficiency consistently (**Supplementary Fig. S5**), which was high enough to adapt to plate reader measurements in a 96-well format.

To evaluate the promoter/terminator activity levels, we used a dual luciferase system with highly sensitive nano luciferase (NLuc) as the reporter and firefly luciferase (FLuc) for normalization^41^. To account for possible batch-to-batch differences in overall protoplast transformation rates/efficiency, we included *FLuc* gene (*UBQ10-FLuc:UBQ5*) in each construct and calculated the NLuc/FLuc ratio as RLU (Relative Light Unit).

For the promoter testing, seventeen promoters drove *NLuc* expression, with termination by either the *NDUFA8* or *HSP* terminator (*Promoter:NLuc:HSP/NDUFA8*). The *UBQ10* promoter exhibited by far the highest expression activity among the promoters, followed by *MAS* (**Fig. 2a,b**). The *HSP* terminator increased gene expression for all promoters except *TUB9*. Two of the newly isolated promoters, *UBQ11* and *UBQ4*, matched or exceeded the activity of the *35S* and *OCS* promoters. Furthermore, the newly isolated promoters *ACT7*, *TUB2*, *TUB9*, *APT1*, *ACT2* and *LEC2* outperformed the commonly used the *NOS* promoter. The *FAD2* and *NDUFA8* promoters had the lowest expression.

**Figure 2.**
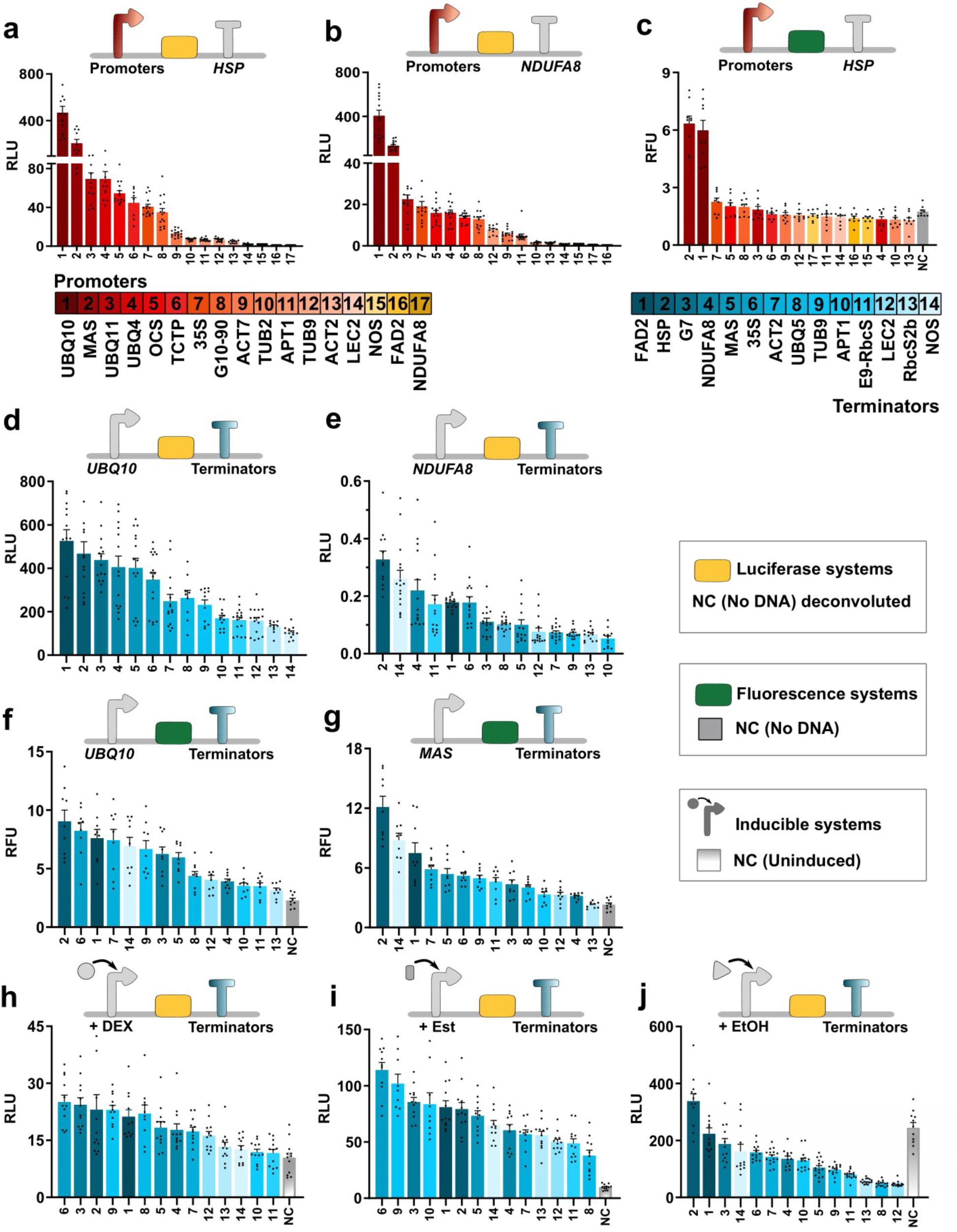
Promoter/terminator part characterization underscores terminator-mediated gene regulation. Promoter characterization is shown on the top panels **(a-c),** the terminator characterization on the middle panels **(d-g**) and inducible systems on the bottom panels **(h-j).** The promoter library (**a-c**) was characterized either with the *HSP* (**a,c**) or *NDUFA8* (**b**) terminator. The terminator library (**d-g**) was examined either with the *UBQ10* (**d,f**), *NDUFA8* (**e**) or *MAS* (**g**) promoters. Inducible systems were induced using **(h)** 2.5μM Dex (*pOp6-35S*), (**i**) 5μM estradiol (*lexA-35S*), **(j)** or 0.1% EtOH (*alcSynth*). Arabidopsis mesophyll protoplasts were transformed with PEG using an optimized transient expression protocol. Relative Light Unit (RLU) refers to the NLuc readout divided by the FLuc readout for **a,b,d,e,h-j**. RFU (Relative Fluorescence Unit) is the sfGFP readout divided by the mScarlet-I readout for **c,f,g**. The bar graphs show RLU or RFU in mean ±SE. The promoter and terminator parts are colour-coded according to the activity strengths in the first characterization series (**a** for promoters; **d** for terminators). The negative control (NC) consisted of untransformed protoplasts (no DNA added). For luciferase systems, NC values were subtracted from measured sample reading during deconvolution, making the NC values 0. For fluorescence samples, deconvolution was not performed, and NC values are displayed. For inducible systems, a first deconvolution round was performed to account for background autofluorescence, and then uninduced *HSP* terminator constructs were used as a control for tightness of gene expression. Data points are from three biological replicates, each with three technical replicates.

For the terminator evaluation, the *NLuc* expression was driven by a strong (*UBQ10*) or weak (*NDUFA8*) promoter and one of the 14 terminators (*UBQ10/NDUFA8:NLuc:Terminators*). The luciferase expression varied by 5.3-6.3 fold for the *UBQ10* and *NDUFA8* promoters, respectively, depending on the terminators they were paired with (**Fig. 2d,e**). For the strong *UBQ10* promoter, the *FAD2* terminator had the highest activity (547.9 RLU), while the *NOS* terminator had the lowest (103.9 RLU). For the weak *NDUFA8* promoter, the *HSP* terminator led to the highest expression (0.327 RLU) and *APT1* to the lowest (0.052 RLU).Since it was surprising to see a wide range of gene expression levels led by different terminators under the same promoter, we extended terminator characterization with the three chemically inducible systems popularly used in plant sciences: dexamethasone (Dex), estradiol, and ethanol inducible systems^42–44^. They are all based on two-component mechanisms involving at least two transcriptional units. The exogenously applied chemical activates the transcription factor that further transactivates the downstream target genes. The target genes are activated by the specific promoters that contain binding sites for the transactivator (*pOp6*, *lexA,* and *alcSynth*), and hence promoters cannot be changed, while terminators can be.

Interestingly, the different terminators resulted in more uniform *NLuc* expression, with a 2.2- and 2.9-fold range in expression levels for the Dex and estradiol inducible promoters (*pOp6-35S* and *lexA-35S*), respectively (**Fig. 2h,i**). For the Dex system, RLU was spread between 37.9 and 111.6 in combination with the *UBQ5* and *35S terminators.* For the estradiol promoter (*lexA*), the *E9-RbcS*and *35S* terminators had RLU counts of 11.6 and 25.1, respectively. In contrast, the ethanol inducible promoter (*alcSynth*), showed a much wider range, with the *HSP* terminator driving sevenfold higher expression than the *LEC2* terminator (Fig. 2j). Both the Dex and estradiol systems showed a basal expression of around 10 RLU; with Dex inducing a ∼11-fold and estradiol a ∼3-fold activation. The ethanol system had high basal expression (i.e., it was leaky), leading to only 30% increase in luminescence upon chemical induction.

### Promoter-coding sequence-terminator interactions in gene regulation

Reflecting on the observed promoter-terminator interactions, we investigated whether changing the coding sequence also influences promoter/terminator activity. Fluorescent proteins are an alternative visible reporting system to luciferases. However, using fluorescent proteins in a plant chassis can be challenging as plants emit red and green-range autofluorescence from their chloroplasts, and stress-induced blue-range autofluorescence from their cytoplasm^45^. Over the years, fluorescent proteins with improved brightness and expression dynamics have been developed, but their quantitative efficacy was not comprehensively characterized in protoplasts. Therefore, we screened fluorescent proteins from four spectrums (green, red, yellow, and blue) to examine their compatibility with a protoplast system using a plate reader or microscopy (**Supplementary Table S4**). Generally, expression of fluorescent proteins was detected 6 hours after transformation and plateaued around 15 hours (**Supplementary Fig. S6**). Informed by this screening, we selected the brightest two fluorescent proteins from different spectra: sfGFP and mScarlet-I. *sfGFP* was the main reporter, while *mScarlet-I* was used as the normalizing gene (similar to *FLuc* above) and expressed using the *UBQ10* promoter and *UBQ5* terminator.

Initially promoters were evaluated with the *HSP* terminator for *sfGFP* expression. Unlike luciferase reporters, not all promoters drove strong enough expression that could be detected with a plate reader (**Fig. 2c**). The highest expression was driven by the *UBQ10* and *MAS* promoters, followed by the *35S* promoter. Readouts from the rest of the promoters could not be distinguished from the background autofluorescence. Therefore, the *UBQ10* and *MAS* promoters were analyzed in combination with all the MAPS terminators **(Fig. 2f,g).** The *HSP* terminator was the strongest with both promoters (9.0 and 12.1 RFU for *UBQ10* and *MAS*, respectively). The *FAD2* terminator was on the high-expression side for both promoters, along with the *UBQ10 promoter-35S terminator* and *MAS promoter-NOS terminator* combinations. The *Rbsc2b* terminator resulted in the lowest expression with both promoters. Overall, a 3-fold expression difference was observed by using different terminators with the *UBQ10* promoter, and the difference was 6-fold with the *MAS* promoter.

Our part characterization revealed that although promoter choice could dominantly determine gene expression strength in some cases (*e.g., pOp6* and *NDUFA8*) (**Fig. 3a**), TU activation by a promoter, terminator or coding sequence is not independent or additive, but combinatorial and even synergistic (**Fig. 3b**). A clear example of such all-part interactions is seen with the *NOS* terminator, which in combination with the *UBQ10* resulted in weak, *NDUFA8* strong, *lexA* medium, *pOp6* weak and *alcSynth* mid-level expression in the luciferase system. When we switched to a fluorescence-based reporter, the combination of the *NOS* terminator with the *UBQ10* and *MAS* promoters drove medium and strong expression, respectively. Among the 14 terminators characterized, only four exhibited stable behaviours with both coding sequences: *FAD2* and *HSP* were consistently on the strong side, while *LEC2* and *RbcS2b* tended towards the weak side.

**Figure 3.**
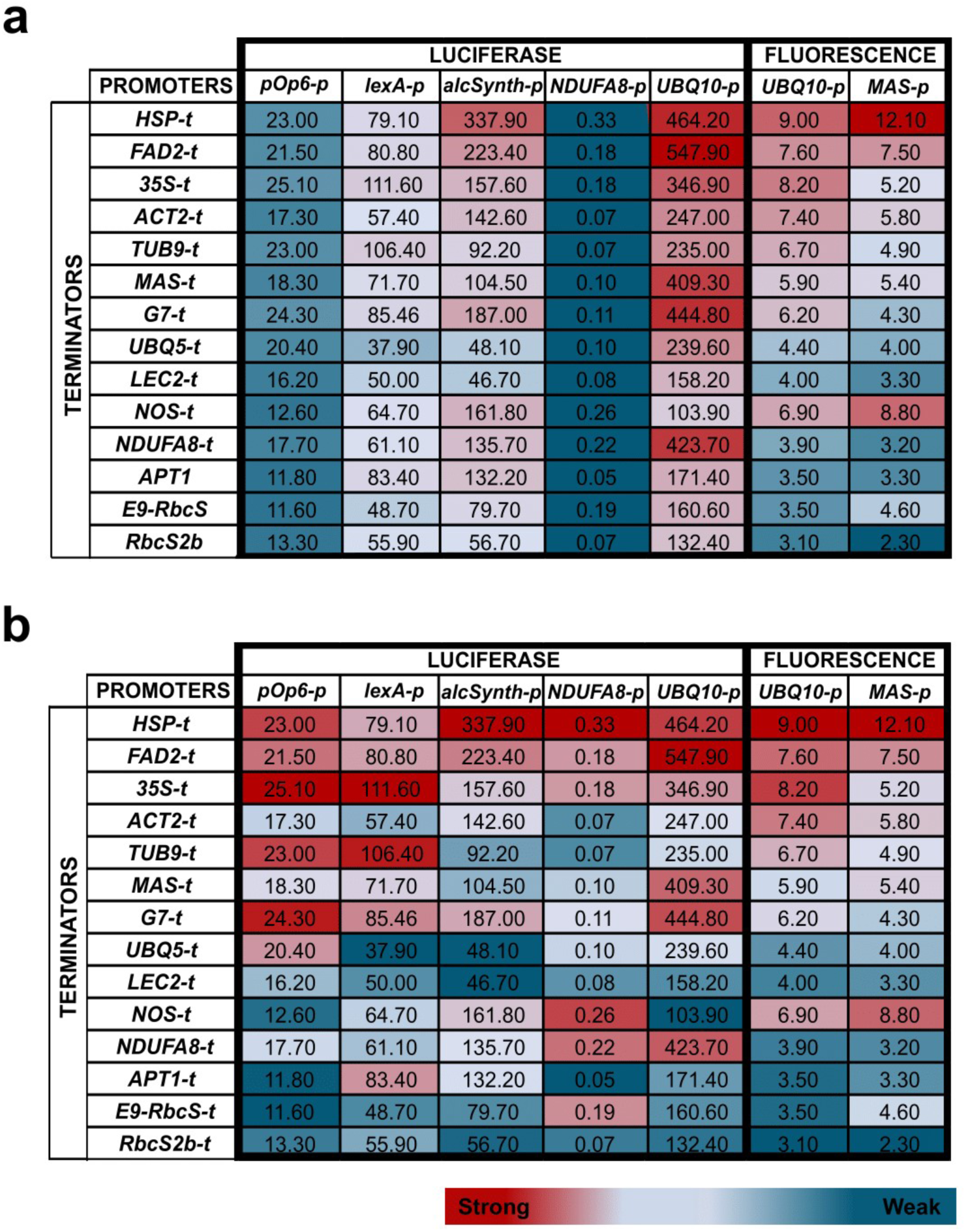
Combinatorial gene regulation by promoters, coding sequences, and terminators. Summary of the reporter expression levels measured for the promoter-coding sequence-terminator combinations. **a.** Heatmap showing all sample comparisons. The reporter gene expression levels were compared across the luciferase samples and the fluorescent protein samples separately. This highlighted that some promoters are dominant and consistent in gene expression strength (*e.g*., *NDUFA8*, *pOp6*). **b.** Heatmap showing the relative strength among the terminators. The color gradient is set independently in each promoter series; dark blue, light blue, and red denote low, mid-level, and high expression, respectively. Dark blue denotes the lowest expression, light blue mid-level, and dark red the highest.

### Dissecting the mechanism of the combinatorial gene regulation

Next, we sought for insights into how the promoters, coding sequences, and terminators interacted in gene regulation. The interactions may regulate gene expression by changing the transcript abundance or with post-transcriptional modifications affecting translation. To distinguish these two possibilities, we performed qPCR to examine how the transcript (mRNA) level correlates with the reporter readout. Because transforming enough protoplasts for RNA extraction is laborious, we selected key constructs to test. The *HSP* and *FAD2* terminators were chosen for consistently strong expression regardless of different promoters and reporter protein sequences. For consistently weak expression, the *NOS* and *Rbcs2b* terminators were selected for *NLuc*, whereas the *Rbcs2b, APT1* and *E9-RbcS* terminators were chosen for *sfGFP*. The *NOS* terminator was chosen because its relative strength varies the most depending on the partnering promoters or coding sequences. Similarly, the *35S* terminator was selected for its variability in strength, although it tends to be on the strong side.

The qPCR assay revealed that the consistently strong *FAD2* and *HSP* terminators have significantly higher mRNA levels; similarly, the weak *E9-Rbcs2b* terminator had significantly lower mRNA levels than the other terminators (**Fig. 4a**). Generally, *NLuc* displayed a linear correlation between the measured reporter expression levels and mRNA levels (**Fig. 4b**), while *sfGFP* exhibited a looser correlation (**Fig. 4c**). Surprisingly, the *35S* terminator yielded higher reporter expression in both reporter combinations compared to its mRNA levels (**Fig. 4b,c**). Taken together, the transcript level analysis suggests that the combinatorial gene regulation is partially explained by transcript abundance and likely to involve post-transcriptional (post-mRNA formation) processes in some constructs. The *35S* terminator for both *NLuc* and *sfGFP* suggested a post-transcriptional enhancer effect, while *NLuc:NOS*, together with *sfGFP:RbcS2b* and *sfGFP:APT1* constructs indicated post-transcriptional repression.

**Figure 4.**
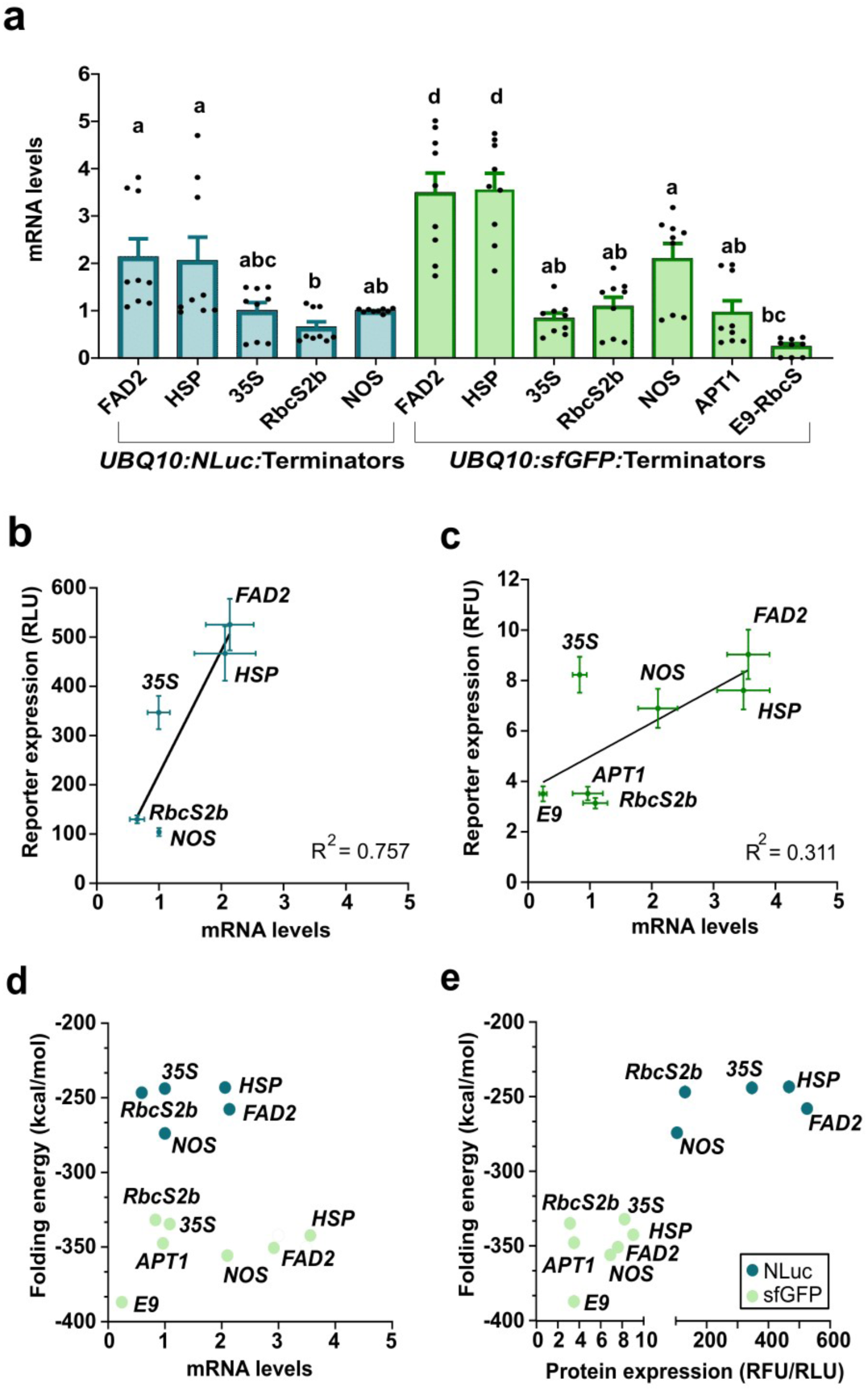
Transcriptional or post-transcriptional regulation underlie combinatorial gene expression control. The mRNA and protein expression levels were compared to differentiate transcriptional or post-translational regulation in different promoter-coding sequence-terminator combinations. **a** Transcriptional regulation: mRNA level was quantitated by qPCR for *NLuc* or *sfGFP* gene expression, under *UBQ10* promoter and different terminators. *UBQ10:FLuc:UBQ5* was used as the normalizing construct for *NLuc*, and *UBQ10:mScarlet-I:UBQ5* for *sfGFP*. One-way ANOVA was applied for the statistical analysis. All the data points from three biological replicates are shown on the graph as dots. **b,c** Correlation of the reporter protein activity to the mRNA levels of different terminator constructs: **b** NLuc and **c** sfGFP. Crossing lines indicate the mean ± SE. **d,c** Correlation of predicted RNA folding energy to **d** mRNA or **e** reporter protein activity level. Blue circlels represent *NLuc* constructs, and green circles *sfGFP* constructs. The folding energy prediction values are available in **Supplementary Table S5.**

Transcriptional regulation is mediated by specific DNA sequences that recruit functional proteins to activate or repress processes from transcriptional initiation to stable mRNA formation. Surprised by how effective terminators are in controlling gene expression, we investigated the nucleotide sequence features possibly influencing terminator strength, such as the GC content and presence of likely functional sequences (*e.g*., the canonical poly-A signal AAUAAA and UGUA motifs) (**Fig. 5, Supplementary Fig. S7**).

**Fig. 5.**
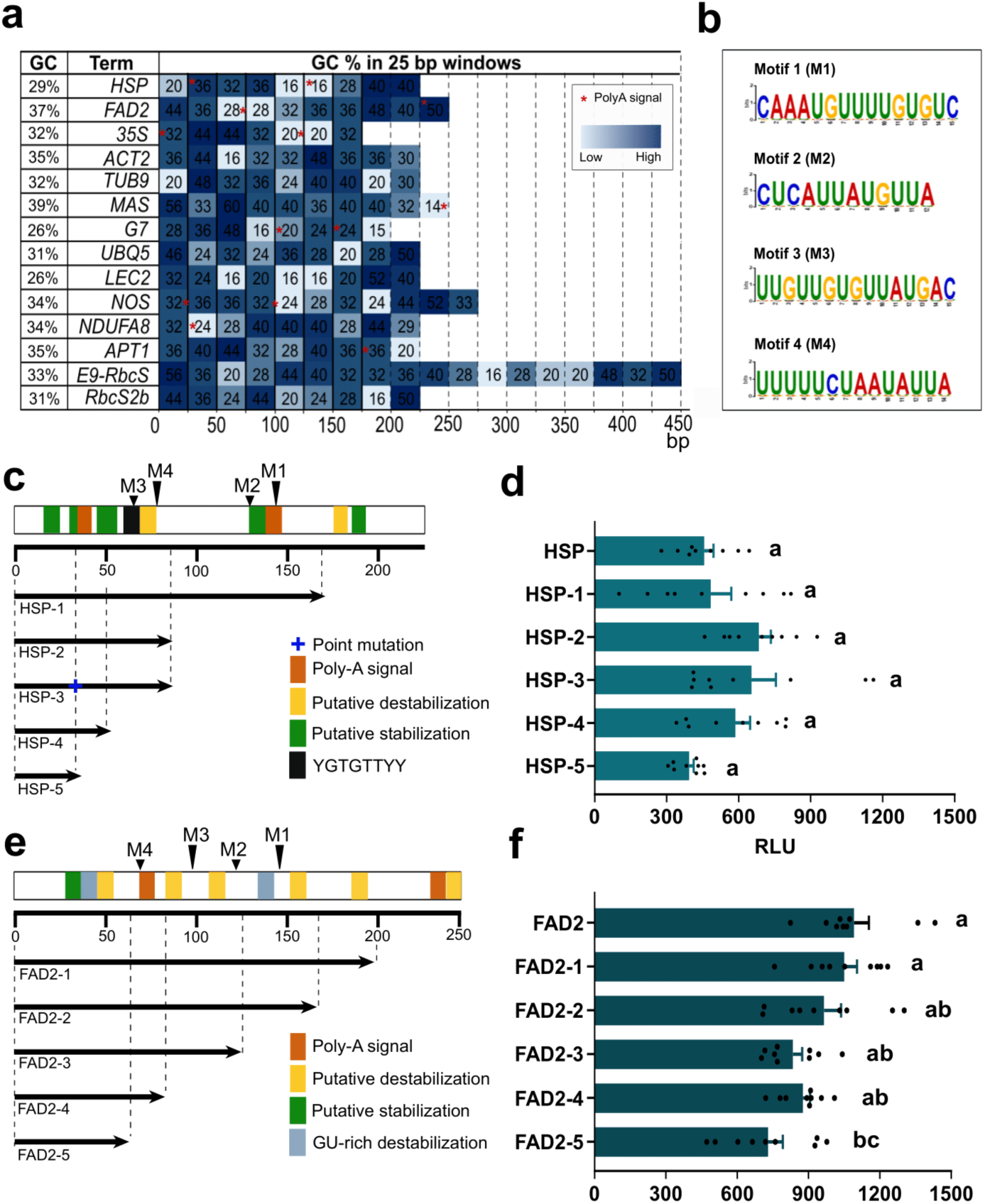
Terminator features that may affect gene expression. **a** GC content of the terminators in the toolkit with 25 bp windows. The shade of the blue indicates the GC content (the darker, the higher). Consistently strong terminators *HSP* and *FAD2*, have a local GC drop of 50-bp that correlates with their dominant Poly-A site (red asterisks). **b** *A priori* identification of motifs in consistently strong terminators. The XSTREME software identified four motifs that are present only in the *HSP-t* and *FAD2-t* terminators. The first identified motif, CAAAUGUUUGUGUC, possibly correlates with high cleave efficiency. **c** Structure, GC content, and functional motifs in the *HSP* and *FAD2* terminators. **c,d.** Deletion series of the *HSP* terminator. **e,f.** Deletion series of the *FAD2* terminator. Schematics of terminator features (**c,e**) and the reporter gene activity (**d,f**) are shown. Relative light unit (RLU) is obtained by normalizing Nluc reporter values by Fluc values. Bar graphs show luciferase activity values in mean ± SE. p < 0.05, one-way ANOVA and Tukey’s HSD test. Data points are from three biological replicates, each with three technical replicates.

A GC content of approximately 30% has been shown to be optimal for synthetic terminator functions, and the average GC content in natural Arabidopsis terminators is about 32.5 ^46^.The GC content in the MAPS terminators had no clear correlations between the strength and GC content of the terminators; consistently or predominantly strong terminators [*HSP* (29.1%), *FAD2* (36.5%), and *35S* (32%)] had a similar range of GC content as weak or variable terminators [*RbcS2b* (30%), *E9-RbcS* (32.8%), *APT1* (35%), and *NOS* (38.7%)] (**Fig. 5a**). One apparent feature in the strong terminators is that the GC content is apparently lower at a 50 bp window, which is located around their dominant Poly-A signal, and then the GC content goes back up. The presence of the canonical AAUAAA poly-A signal tends to enhance terminator strength^46^.

We also searched for sequence motifs that may link to the consistently high strength of *HSP* and *FAD2* terminators. The presence of the UGUA motif around 30-40 bp upstream of the RNA cleavage site is thought to enhance the cleavage and thus increase terminator strength in the *35S* terminator ^46^. The *HSP* terminator has two UGUA motifs at locations 119 bp and 206 bp, while *FAD2* lacks the motif. Interestingly, the XSTREME motif discovery tool identified four motifs that putatively have functional roles *a piriori* (**Fig. 5b,c,e**). One of them is the 15-bp motif (CAAAUGUUUUGUGUC) found around 145 bp in both the *HSP* and *FAD2* terminators, which correspond to the transcript cleavage site. Three other motifs - CUCAUUAUGUUA, UUGUUGUGUUAUGAC, and UUUUUCUAAUAUUA - were found at similar locations in both terminators but around 10-20 bp apart. Only the CUCAUUAUGUUA motif is present in the *UBQ5* terminator at 167 bp (**Fig. 5b**). The effects of the UUUUU motif are complicated; it decreases terminator strength in maize protoplasts, especially if they surround a UGUA motif, while U-rich sequences increase terminator strength in tobacco leaves^46^. Many of these AT/U-rich motifs reside inside the 50 bp low GC domains described above.

We examined similar potentially functional features in the ‘outlier terminators’ – the terminators performing stronger or weaker than expected for their transcript levels (*i.e*., likely post-transcriptionally regulated) (**Fig. 4a,b**). The GC content of outlier terminators is moderate and varies between 30.5-35.0% (**Fig. 5a**). *RbcS2b* is the only outlier terminator that does not possess the canonical poly-A signal. The *35S* terminator has three repeats of the UGUA motif (x3 starting from 96 bp separated by UU). The UUUUU motif is present in the *NOS*, *RbcS2b* and *APT1* terminators **(Supplementary Fig. S6).** Therefore, no distinct sequence signatures were identified to differentiate the terminators that are primarily regulated transcriptionally from the outliers.

We then wanted to experimentally probe how the above-identified sequence features influence the terminator activity by generating a deletion series of the *HSP* and *FAD2* terminators. Five terminator variations were made to sequentially delete possible regulatory elements, such as Poly-A signals, putative destabilization signals, and Musashi binding elements (**Fig. 5c-f**). Statistical analysis of the results showed only a significant difference between the full-length terminator (240 bp) and *FAD2* Sequence 5, which is the shortest variation (70 bp) with no Poly-A site, as well as the deletion series Sequence 1 (200 bp). The rest of the *HSP* series, as well as the *FAD2* series, showed no statistically significant difference in reporter expression (**Fig. 5c-f**). This result suggests short sequences (30-50 bp) at the 5’ end of terminators might determine the gene expression strength, and terminator functional elements remain unresolved, especially in plant contexts.

We also investigated a sequence feature likely enhancing promoter activity. *UBQ10* is by far the strongest promoter in the MAPS toolkit (**Fig. 2,3**), and its TSS structure is predicted to be unstructured and single-stranded (**Fig. 6**). In the survey of the Arabidopsis genome, translation efficiency was found higher for transcripts with unstructured 5’UTR^47^. To test if the strength of the *UBQ10* promoter is dependent on the open loop structure of its 3’ UTR, which consists of mostly TSS, three mutated versions were created. The first version, Mutated Control (MUTC), involves point mutations that preserve the predicted loop structure; therefore, it is expected to behave similar to the original *UBQ10* TSS sequence. Additionally, two other versions, a tight stem-loop structure Mutated 1 (MUT1) and a slightly more branched but more loosely stemmed Mutated 2 (MUT2), were designed with point mutations (**Fig. 6a,b**). TSSPlant and Softberry software were used to confirm the mutations do not interfere with the identified motifs or introduce new motifs compared to the WT.

**Figure 6.**
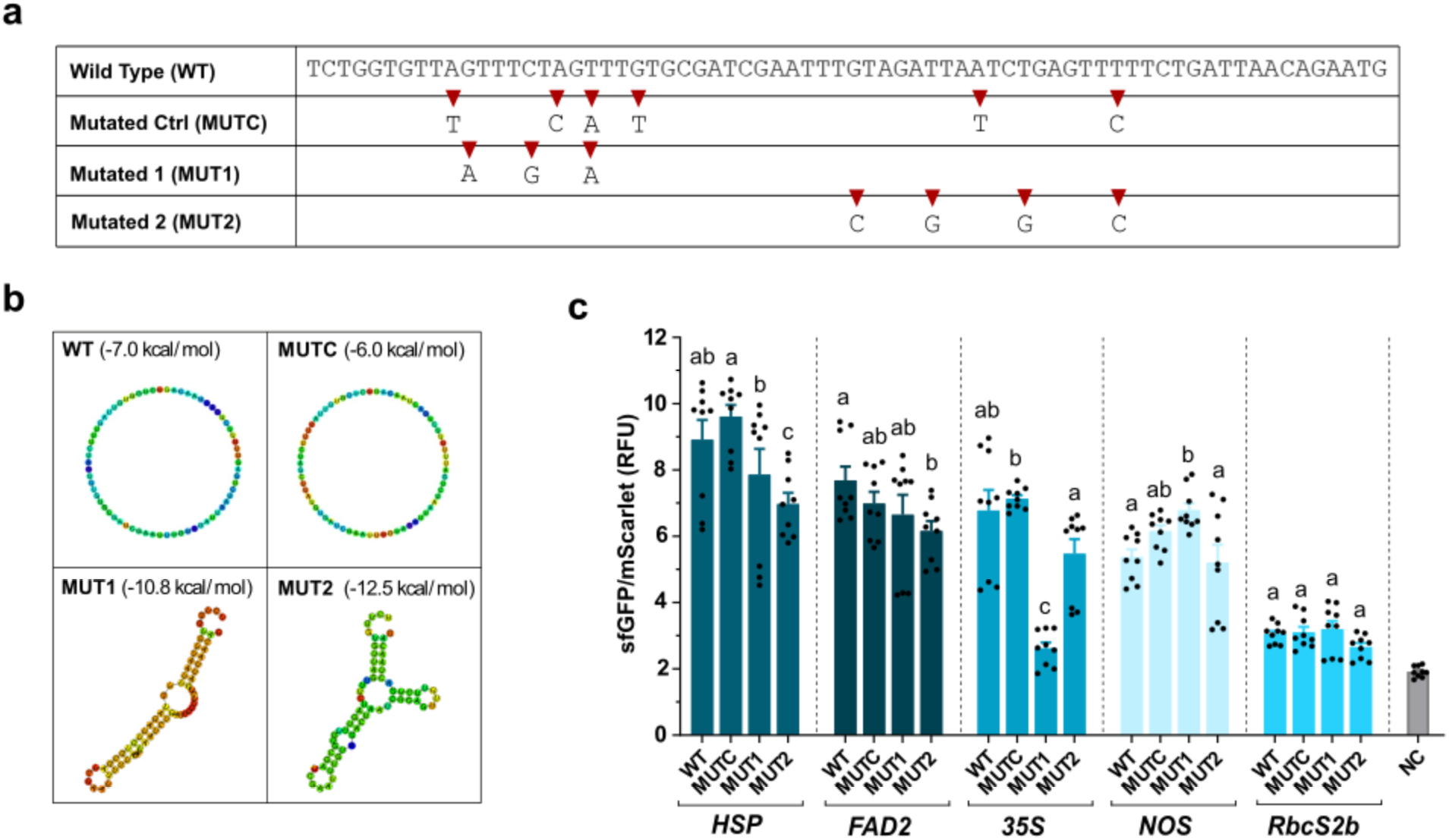
Open structure at *UBQ10* promoter TSS affects gene expression. Strong promoters, such as *UBQ10*, tend to have an open loop structure at the TSS. To test the significance of this structural signature, point mutations were introduced to form closed stem-loop structures instead. **a** Four TSS variants were tested: Wild Type (WT), Mutated control (MUTC), Mutated 1 (MUT1), and Mutated 2 (MUT2), and **b** their predicted RNA folding structures. **c** Open loop structure at TSS enhances gene expression. *sfGFP* was expressed under *UBQ10* containing WT, MUTC, MUT1, or MUT2 variant of TSS. Five different terminators were chosen to assess the effect: *HSP, FAD2, 35S, NOS,* and *RbcS2b*. The normalization construct was *UBQ10:mScarlet-I:UBQ5* for all samples. Bar graphs show mean reporter activity values ± SE. Negative control (NC) was untransformed protoplasts. One-way ANOVA with Tukey’s HSD was done individually for each terminator category.

When the TSS variants were used to drive *sfGFP* with five different terminators (*HSP, FAD2, 35S, NOS,* and *RbcS2b*), the results revealed no statistically significant difference in expression between WT and MUTC, as expected (**Fig. 6c**). However, MUT2 exhibited a reduction in expression by approximately 20% for the *HSP* and *FAD2* terminators. Additionally, MUT1 resulted in a 60% decrease in expression with *35S*, while resulting in a 30% increase with the *NOS* terminator. There was no difference between the three mutated versions and the WT for the *RbcS2b* samples, where the WT expression is already very weak (**Fig. 6c**). With the strong terminators, reporter expression was reduced in MUT1 and MUT2 compared to WT and MUTC variants, suggesting that conversion from open loop to stem-loop decreases gene expression.

To gain insights into how promoters, coding sequences, and terminators interact post-transcriptionally, we studied RNA folding. RNA is single-stranded and extensively forms secondary and tertiary structures via hydrogen bonds bringing together (nearly) complementary sequences. Such 2D and 3D structures (*e.g*., G-quadruplex) can strongly influence translation and protein expression^48,49^. The combination-dependent regulatory function among the three TU parts may be explained by direct physical interactions through RNA folding.

Using the RNAFold software^50^, the folding energy of the whole mRNA sequence was calculated for the transcript species we selected for the qPCR analysis above. The transcription start site (TSS) was identified based on the TSSPlant software^51^, and the downstream promoter sequence was incorporated, along with the protein-coding and terminator sequences. The lower the holding energy, the tighter the transcript folds, and the less likely for translation to occur. No direct correlation was found between the predicted transcript folding energy and mRNA expression levels or between the folding energies and reporter gene expression (**Fig. 4d,e**). We therefore proceeded to examine the RNA folding structure and local interactions among the nucleotide sequences.

We then visually examined the RNA folding structures in 2D. RNA secondary structure formation was predicted using the ViennaFold2.0 software^50^, in which the whole transcript (identified as described above) was used as the input. Within the transcript, the strong *UBQ10* and *MAS* promoters are likely to have little interaction with the other parts: no interaction with the *HSP* and *RbcS2b* terminators and minimal to medium interactions with the *FAD2* terminator were predicted by the software (**Fig. 7**). On the contrary, *NOS,* a highly variable terminator, may have strong interactions with the other parts (*UBQ10:NLuc*) when it drives weak reporter expression. Interestingly, there was no apparent correlation between inducible promoter-terminator structures and reporter output, except for *pOp6:NLuc:NOS* (**Supplementary Fig. S8**). In strong and consistent terminators like *HSP*, we tended to find loops bigger than 20 bp **(Fig. 7).** When the variable strength-terminator *NOS* is paired with strong promoters (*UBQ10* and *MAS),* it also may form a large loop structure, though not when combined with other promoters (*e.g*., *NDUFA8*) (**Fig. 6, Supplementary Fig. S9**). RNA sequence-mediated cross-part interactions among the promoters, coding sequences, and terminators could explain the combinatorial gene regulation.

**Figure 7.**
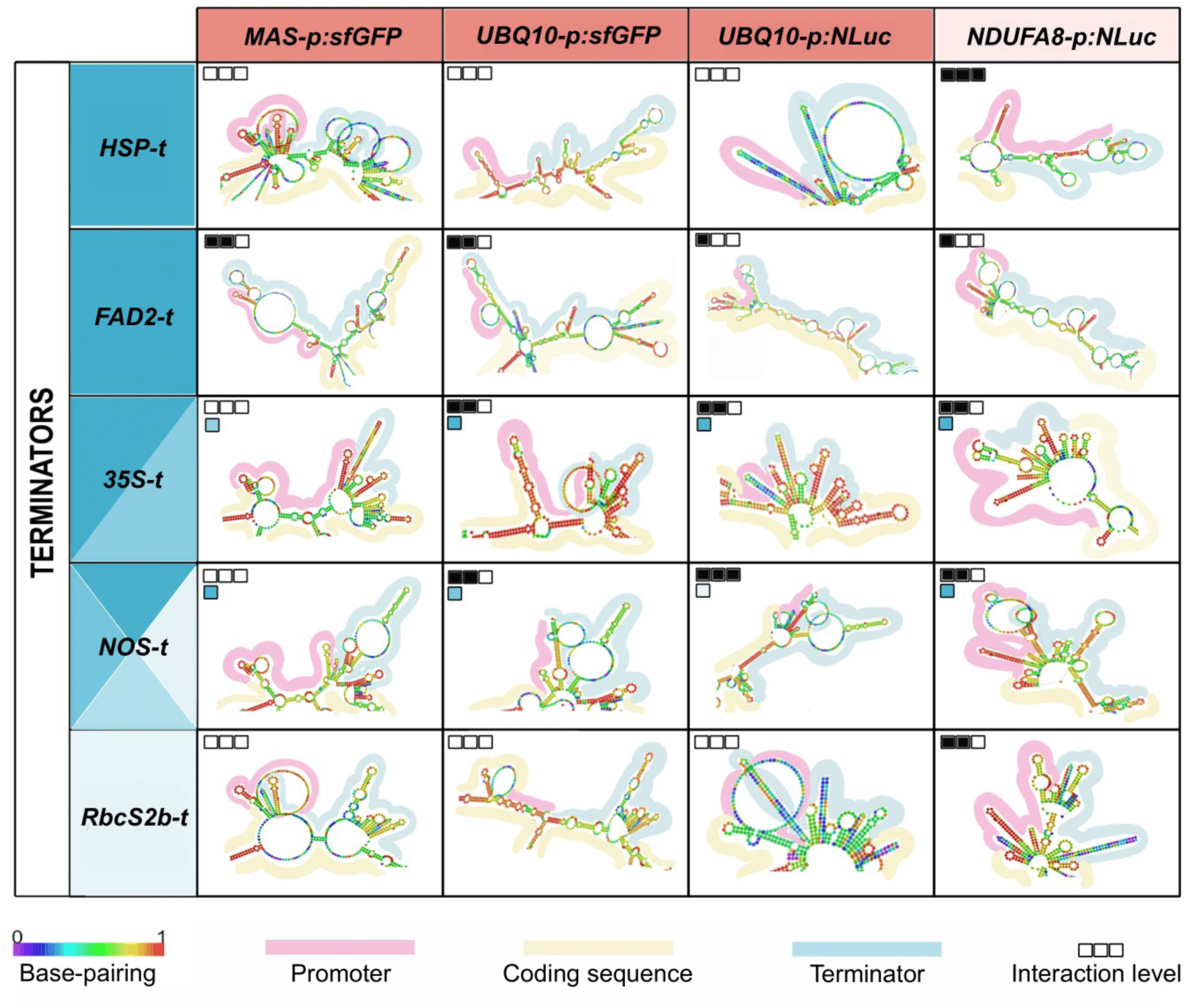
Structural interaction patterns in the transcript RNA sequences. RNA folding was visualized to show the likelihood of intra-transcript nucleotide interactions. The promoter part corresponds to the TSS of the *MAS, UBQ10* and *NDUFA8* promoters. Terminator strength is represented as a blue gradient, where dark blue represents strong and light blue weak; terminators whose strength shifts depending on the promoters are marked with mixed shades. Promoter strength is represented similarly, where dark to light red corresponds to strong to weak. Interaction levels are categorized as none (empty squares), weak (one square full), medium (two squares full), and strong (three squares full). For none, there is no base pairing predicted between promoter and terminator parts; for weak, there is only base pairing with blue and green colours, which means it is <0.5 probable; for medium, there is base pairing in yellow and orange colours, which means 0.75 probability, or 5 bp with red base pairing, indicating ∼1.0 probability. Strong interactions are defined as >5 red base pairs.

## DISCUSSION

### MAPS: A new synthetic biology toolkit for engineering with plant systems

In the present work, we established the Mobius Assembly for Plant Systems (MAPS), which is an adaptation of the Mobius Assembly^26^ for plant genetic programming (**Supplemental Figure S1, Supplemental Table S1**). MAPS is a stand-alone yet highly versatile and functional DNA assembly platform and genetic toolkit collection. It is based on small binary vectors for all levels of cloning and comes with a well-characterized, Phytobrick-compatible standard part library for gene expression control in plant species, including a suite of reporter protein cassettes. We have also introduced a new feature for Mobius Assembly to facilitate the generation of a series of constructs with part variations; MethyAble exploits *in vitro* DNA methylation to directly feed Phytobricks in single- or multi-TU constructs.

For plant genetic engineering, small binary vectors are instrumental in enhancing efficiency in cloning and transformation. As we wanted a reliable core vector in our kit forboth cloning and transformation applications, we developed a new binary vector pMAP. This new vector has a unique dual-mode origin of replications for *E. coli* and *Rhizobium* backgrounds, which enables amplification in high copy numbers for experiments demanding high DNA yields, such as the protoplast assay (4 ug DNA per 75 ul transformation) used in this study. The plasmid dimerization phenomenon we have observed is not unique to Mobius Assembly vectors; in fact, with the increasing availability of whole-plasmid sequencing technologies, more dimers are being detected. Therefore, including a *cer* site in the backbone provides an effective method for preventing dimerization and growth defects (**Supplementary Fig. S3**)^38^.

MAPS delivers a collection of short promoters (17 constitutive and three inducible) and terminator (14 constitutive) standard parts for plant expression. Ten of the promoters and six of the terminators were newly isolated, and they are short in length: 300-600 bp (promoters) and 200bp (terminators). We confirmed the short size of the promoters did not compromise their strength. The *Actin 2* promoter characterized in MoClo and GB2.0 toolboxes had comparable activity to the NOS promoter^52,53^; the shorter version in the MAPS toolkit, *ACT2* (340bp) and *NOS* promoter, exhibited similar expression activity (**Fig. 3**). In general, the size of the available standard promoter or terminator parts for plant engineering (*e.g*., MoClo plants, GB plants, and GreenGate) can be as long as 4 kb. Shorter parts are desirable as large constructs can reduce plasmid stability in bacteria, while also lowering transformation efficiency and causing incomplete/truncated transformation in planta^14,15^.

Here, we reported the first characterization of the MAPS toolkit, which was done exclusively in Arabidopsis leaf mesophyll protoplasts with a plate reader-based high-throughput assay. Most plant standard part characterization to date has been conducted in either Arabidopsis protoplasts or *Nicotiana benthamiana* (leaf expression system), which are both dicot plants. The standard parts can behave differently in different backgrounds. In monocots, for example, the *35S* and *NOS* promoters drive low activity in monocot *Setaria viridis*^18,54^. Even within dicots, the same part can have different levels of expression^52,53^. The *OCS* promoter had strong activity in our study but low expression in *N. benthamiana*^52^. Genetic parts can behave differently in different chassis, and we should not generalize characterization results among different plant species or expression systems (*e.g.*, whole plants, leaves, protoplasts, or cell cultures). We recommend not making assumptions and testing the activity levels of standard parts in the specific context in which users want to apply them.

### Terminators influence gene expression, in part through interaction with promoters

Our characterization of the MAPS terminators underscored the strong influence of the terminators cast on gene expression. It is a common practice to rely on promoters to control gene expression level and overlook the impact of terminators. For example, when the domestication of the synthetic plant promoter *G10-90* promoter resulted in loss of expression, researchers replaced it withthe *AtRPL37aC* promoter to rectify^44^. They were using the *psE9-RbcS* terminator, which in our assay had low activity; the gene expression could potentially have been restored with a stronger terminator instead. Even though promoters generally have a strong influence on gene expression (**Fig. 3a**), terminators could alter the gene expression up to 8-fold in this study (**Fig. 2,3**). Alternative to constitutive expression, chemically inducible systems are powerful tools to control the timing, localization, and amplitude of transgene expression. Dex-, estradiol-, and ethanol-inducible systems were developed 20 years ago and have been used in numerous studies since then^42^. The extent of target gene induction can be modulated by exchanging the terminators (**Fig. 3**).

Terminators influence gene expression through multiple mechanisms^55^, including post-transcriptional regulation via the 3’ UTR, which can affect mRNA stability through specific sequence motifs. For instance, certain motifs at the 3’ UTR in Arabidopsis can stabilize (e.g., TTGCTT) or destabilize (e.g., AATTTT) mRNA^56^. Poly(A) tails also play a role, with short tails often linked to strong gene expression ^57^. In Arabidopsis, the poly(A) tail can prevent RDR6 from converting aberrant mRNAs into degradation substrates^57^. Unpolyadenylated transcripts derived from terminator-less constructs or readthrough mRNAs from transgenes with strong promoters are subjected to RDR6-mediated silencing^59^, which might explain the increased gene expression observed with double terminators^60–62^. However, the synergistic effects of terminators, particularly when combined with weak promoters, suggest interactions beyond these mechanisms. Direct interactions between terminators and promoters, potentially involving gene looping mediated by DNA-binding proteins, can influence gene expression^63^. These interactions might affect transcriptional memory^64^, intron-mediated modulation of transcription^65^, transcription directionality^66^, reinitiation of transcription^67^, and transcription termination^68,69^. Additionally, terminators may impact epigenetic regulation, as the absence of a terminator can lead to increased DNA methylation on promoter regions, silencing transgene expression^70^.

### Promoter-coding sequence-terminator interactions in gene regulation

In addition to the interactions between terminators and promoters, we observed that protein-coding sequences can also modify the strength of promoter-terminator function. The coding sequence may impact gene expression through factors such as codon usage bias, gene length, GC content, correct 5’ cap, and mRNA folding^71^. Both the *NLuc* (516 bp, GC 52.7%) and *sfGFP* (714 bp, GC 61.3%) have been optimized for plant expression in terms of codon usage. In an orthogonal system where part performance does not depend on the context, any factors related to codon usage, gene length, GC content, or correct 5’ cap may affect the magnitudes of the reporter readout, but the comparative profiles of gene expression across the promoter-terminator pairs should remain the same. However, the relative strengths of promoters and terminators changed when the reporter protein was replaced (**Fig. 3,4**), indicating that the coding sequence interacted with the promoter and the terminator presumably directly and physically.

We wanted to understand how the promoter-coding sequence-terminator interactions occur. There are many possible mechanisms (some examples outlined above), which likely to influence gene activity in combinations. However, to simplify, such interactions can be explained either at the DNA or RNA level – or transcriptional or post-transcriptional mechanisms. The qPCR assay indicated that the reporter genes activity was regulated transcriptionally by controlling the stable transcript abundance, but also post-transcriptionally as the transcript abundance alone did not always correlate with the reporter activity (**Fig. 4**).

To explore the mechanisms behind the transcriptional regulation, we surveyed DNA sequence features potentially involved in determining terminator or promoter activity. The terminators with consistently strong reporter protein readout (*HSP* and *FAD2*) also had high mRNA levels (**Fig. 5a**). These two terminators contain a local drop (50 bp) in the GC content, that was absent in most other terminators. It is likely that this low GC region holds other structural features that aid the recognition of transcription termination and cleavage. When other possible motifs were searched for *a priori* in *HSP* and *FAD2 but not in other MAPS terminators,* the XSTREME motif discovery tool identified four motifs. Among the four, CAAAUGUUUGUGUC motif at location 145 bp in *HSP* and 146 bp in *FAD2*, may aid with cleavage site recognition and cleavage efficiency, enhancing overall terminator strength (**Fig 7b,c**).

With our deletion series, we have seen that only the shortest FAD2 terminator (70 bp) has significantly reduced strength compared to the full-length version (240 bp). The first 35 bp of the *HSP* terminator has a strength statistically nondifferent compared to the full-length (**Fig. 5c,d**); this is consistent with Felippes *et al*.^72^, which showed that the first 32 bp of the *HSP* terminator is a transferable element that enhances the strength of other terminators. A very short (<50 bp), minimal terminator domain may govern much of the terminator activity, especially in transient expression systems, whereas the specific features identified (**Fig. 5**) may play clearer roles in longer timescales or certain physiological or developmental contexts.

Recent studies revealed the roles of RNA structures in gene expression control. Lower mRNA folding energies, typically associated with greater structural stability, impact ribosome function and translation in organisms such as *E. coli* and *S. cerevisiae*^73,74^. Beyond global folding, local secondary structures can also play a pivotal role in shaping gene expression. Pseudoknots, hairpins, loops, and the presence of hydrogen bonding patterns upstream of the transcription start site have been shown to reduce ribosome sequestration and translation rates^75,76^. A study with 224,000 synthetic sequences in *E. coli* has highlighted the importance of secondary structures at the 30 bp region upstream and downstream of the start codon (referred to as STR-_30:+30_)^77^. While comprehensive studies in plants are still pending, *in vivo* genome-wide profiling of Arabidopsis RNA secondary structures revealed that unstructured regions upstream of the start codon are enriched in high translation efficiency mRNAs^47^. In the strong promoter *UBQ10*, we found an unstructured TSS (loop) where altering its structure affected reporter output (**Fig. 6**).

RNA secondary structures also point to a similar regulatory influence in terminator-dependent gene regulation. When the consistently strong terminators *HSP and FAD2* were combined with various promoters, loops of >20 bp were predicted (**Fig. 7**). This might be a structural feature of strong-acting terminators, since a similar >20 bp loop was predicted when the variable *NOS* terminator was combined with strong promoters (*MAS, UBQ10);* yet the loop is thought to re-configure to a stem-loop structure when combined with weaker promoters. Expanding on these structural themes, we examined the folding structure of the terminators from Felippes *et al*.^71^ that showed increased strength when combined with the first 32 bp of the *HSP* terminator. The predicted folding of the transcript showed that the addition of the 32 bp led to an increase in the size of the loops >20 bp for the *NOS* and *ACS2* terminators, as well as higher base-pairing probability structures for *Rbcs1A* (**Supplementary Fig. S9**).

RNA folding patterns predicted the consistent activity promoter and terminator parts tend to be isolated in terms of secondary structure, displaying a low level of physical interactions from the other parts, regardless of their sequences (**Fig. 7, Supplementary Fig. S8**). Additionally, in the case of the *NOS* terminator, cross-part interactions could explain the high variability. It is plausible that certain promoters and terminators possess currently unknown regulatory elements that enhance their activity while physically restricting their interactions with other parts. Meanwhile, there may be TU parts that may require physical interactions with other parts to function effectively. For example, in mammalian systems, terminator elements require specific secondary structures to interact between the polyadenylation complex and mRNA^78^. Synthetic parts with known elements, like the *pOp6* and *LexA* inducible promoters, likely hold key to context-independency. Even though their transcripts were predicted to fold, generating cross-fold interactions, such interaction did not seem to have a strong effect on the reporter output (**Supplementary Fig. S8**).

## CONCLUSION

We developed a new, simple and versatile synthetic biology toolkit for plant genetic engineering, Mobius Assembly for Plant Systems. In so doing, we unravelled new insights into gene regulation using a bottom-up approach and have shown that gene expression is regulated by all parts of a transcriptional unit combinatorically. Uncovering the relationship between the structure and function of RNA (and DNA) will help establish the principles for designing consistent and predictive standard parts that are independent of the nucleotide context-independent and more orthogonal.

## MATERIALS AND METHODS

### Bacterial strains and growth conditions

*E. coli* strains JM109, and NEB stable were used*. E. coli* chemically competent cells were prepared in-house using the TSS preparation described by Chung & Miller^79^. *E. coli* cells were incubated at 37°C (30°C for NEB stable), 200 min-1 either in 5 ml (for a high copy) or 10 ml (for a low copy), or 100 ml (for midi prep) in LB growth medium supplemented with antibiotics. Cells bearing the LhGR-pOp6 and sXVE-lexA inducible systems were grown for 24 h instead due to their slower growth rate.

### Bacterial transformations and plasmid isolation

For *E. coli* transformation, 5 μl of the plasmid DNA was incubated with 100 μl of the competent cells on ice for 30 min, followed by a heat shock at 42°C for 90 sec (30 sec for TOP10) and re-cooled on ice for 5 min. S.O.C medium (400 μl) was added, and after 1 h incubation at 37°C, 100 μl of the cell suspension was plated on LB agar plates with antibiotic selection. Plasmids were isolated using Monarch (NEB) Plasmid Miniprep Kits. The GeneJET Plasmid Midiprep Kit (ThermoFisher) were used for higher yields.

### MAPS vector construction

MAPS Level 1 and Level 2 Acceptor Vectors were built using Gibson Assembly. The Mobius Assembly cassettes were amplified from the corresponding *E. coli* plasmids in the Mobius Assembly Vector toolkit^26^ and fused to the plant binary vectors. The pGreen-based Level 1 vectors were constructed using pGreen0029^31^. The *NptI* gene was replaced with the spectinomycin resistance gene amplified from the pCR8 vector (ThermoFisher) to generate Level 2 vectors. The pLX-based Level 1 and Level 2 vectors were built using pLX-B3 and pLX-B2^33^, respectively.

WKS1 Ori was synthesized (Twist Bioscience) into two parts due to repetitive sequences, and pUC Ori was amplified from pUC19, both flanked by *BsaI* recognition sites. The minimum sequence requirement of pUC for stable replication in E. coli was found to include RNA I/RNA II transcripts on the 5′-side and dnaA/dnaA′ boxes on the 3′-side, while co-directional transcription of two different replicons in the same plasmid was shown to increase transformation efficiency and DNA yield^32^. A pLX Level 1 A vector with the construct NOS:BglR:NOS-UBQ10:nluc:HSP was amplified outside the BBR1 Ori with primers harbouring *BsaI* recognition sites and fused with pUC and WKS1 Ori. The resulting plasmid was used as the template to amplify the pUC-WKS1 fused Ori, which then replaced BBR1 Ori from the pLX-based Level 1A and Level 2A vectors with Gibson Assembly, resulting in the pMAP Level 1A and Level 2A vectors. The rest of the pMAP vectors were constructed again using isothermal assembly and as a template for the Mobius cloning cassettes, the plasmids from the original Mobius Assembly kit^26^.

MethylAble modules were devised with the Gibson Assembly using mUAV as a template. Overlapping primers bearing two outward-facing *BsaI* sites prone to CpG methylation and suitable standard overhangs were used to amplify mUAV into two parts. The resulting parts were purified and digested with *DpnI* (ThermoFisher) to eliminate the template DNA, and they were fused in an isothermal reaction.

### Mobius Assembly and standard part library construction

A detailed protocol on Mobius Assembly can be found in^80^. Briefly, the assembly was performed in a one-tube reaction with a total volume of 10 μl, with ∼50 ng Acceptor Vectors and double amounts of inserts. Reagents added were 1 μl of 1 mg/ml BSA (diluted from 20 mg/ml - NEB), 1 μl T4 DNA ligase buffer (ThermoFisher/NEB), 0.5 μl *AarI/PaqCI* (ThermoFisher/NEB) and 0.2 μl 50x oligos of the *AarI* recognition site for Level 0 and Level 2 cloning or Eco31I/BsaI-HFv2 (ThermoFisher/NEB) for Level 1 cloning, and 0.5 μl T4 DNA ligase (ThermoFisher/NEB). The reactions were incubated in a thermocycler for 5-10 cycles of 5 min at 37°C and 10 min at 16°C, followed by 5 min digestion at 37°C and 5 min deactivation at 80°C (5 cycles for Level 0 and the first round of Level 1 cloning – 10 cycles for Level 2 and large constructs >10 kb).

### PCR amplification

All PCR amplifications for plasmid construction and cloning were performed using Q5® High-Fidelity DNA Polymerase (NEB), followed by purification with Monarch® PCR & DNA Cleanup Kit (NEB). Successful DNA assembly was verified first by colony PCR using GoTaq® Green Master Mix (Promega) and then with double restriction digestion with EcoRI-HF (NEB) and PstI-HF (NEB). The constructs were further verified by Sanger sequencing (GATC Biotech-Eurofins, Edinburgh Genomics and Source BioScience) and whole plasmid sequencing (Full Circle).

### Promoter and terminator part design and generation

Likely constitutive promoters and terminators were selected from the literature, especially the genes commonly used as positive controls in qRT-PCR^81^. Their sequences were retrieved from the TAIR webpage^82^ (http://www.arabidopsis.org/index.jsp) and blasted in NCBI to find the untranslated regions flanking the genes. A 1.5 kb sequence upstream of the start codon was run through the online prediction software and TSSPlant^51^ (http://www.softberry.com) to identify TATA and TATA-less promoters or enhancer sites. In the promoter selection, it was also considered, when possible, to include the initiator (INR) elements (YYA(+1)NWYY-TYA(+1)YYN-TYA(+1)GGG)) and downstream promoter (DPE) element (RGWYV). Potential promoter elements linked to increased gene expression were identified using PlantCare^83^ and PLACE^84^ software. Terminators were selected with PASPA, a web server for poly(A) site prediction in plants and algae^85^ (http://bmi.xmu.edu.cn/paspa/interface/run_PASPA.php). A sequence 300bp downstream of the stop codon was input into PASPA, and the end of the terminator was set at 10bp after the second polyadenylation site, resulting in a sequence of around 200bp. They were then analyzed in RegRNA2.0 to identify other RNA functional motifs^86^ (http://regrna2.mbc.nctu.edu.tw).

The parts designed in the first round were short promoters (∼300bp) and short terminators (∼200bp) from the genes *ACT2, FAD2, TUB9, APT1, NDUFA8* and *LEC2.* As the ∼300 bp promoters were very low in activity (Data not shown), we designed a new set of promoters derived from the genes *TUB2, UBQ11, UBQ4, ACT7*, and the longer versions of the previous promoters (∼500 kb). Appropriate primers compiling to the Phytobrick standard overhangs were designed for both promoters and terminators for cloning into mUAV.

### Protoplast isolation and transformation

Arabidopsis (Wildtype, Col-0) seeds were sown on the soil. After a 2-day pre-treatment at 4°C in darkness, they were sawn and grown under long-day conditions (21°C; 16h light / 8h dark cycles; light intensity ∼100μmol/m2s-1; humidity 40-65%) until harvest. The protoplast isolation and transformation protocol was developed based on Chupeau *et al*., (2013)^40^ and Faraco *et al*., (2011)^87^. The optimized protocol is descripted in Supplementary Information, with representative images of consistently high transformation (**Supplementary Fig. S4**).

### Luciferase assay

The 96-well plates containing the transformed protoplasts were let to sediment and 60 μl of supernatant was discarded. The protoplasts were resuspended, and 40 μl was transferred to white optical plates in a grid pattern with empty spaces between wells to reduce luminescence bleed-through. Luciferase activity was assayed in an Omega luminescence plate-reader (Fluostar) with four different gains following the instructions of the Nano Dual-Luciferase® Reporter kit (Promega, N1620). A further correction for luminescence bleed-through and background signal was applied using the software developed by Mauri et al.^88^ Then NLuc signal was divided by the FLuc signal to normalize for the transformation efficiency.

### Fluorescence assay

For fluorescent protein assays, TECAN Spark plate reader was used with either 96-well plates (Greiner) or black Thermo Scientific Nunc F96 MicroWell. Excitation and emission for the super-folder green fluorescent protein (sfGFP) was 485 nm and 520 nm, respectively. For mScarlet-I, excitation and emission wavelengths were 560nm and 620nm. Measurements were taken at two different gains to ensure signals don’t overshoot beyond saturation. The sfGFP values were divided by the mScarlet-I values to normalize for the transformation efficiency. Since fluorescence was not always active, unlike luminescence and also because black-walled plates were used to prevent signal bleed-through, deconvolution correction was not applied to the fluorescent samples.^88^

### RNA isolation and real-time PCR (qRT-PCR)

Isolation and transformation were performed as described above, with reaction and solution volumes upscaled by 26 times. 24 h after transformation, the supernatant was discarded, and sedimented protoplasts were harvested for RNA extraction with GeneJET Plant RNA Purification Kit (ThermoFisher). Extracted RNA was treated with DNA-free DNA Removal Kit (Invitrogen). Then, reverse transcription was performed with UltraScript 2.0 cDNA Synthesis Kit (PCR Biosystems). This cDNA synthesis kit uses optimum amounts of oligo (dT) and random hexamers for unbiased amplification of RNA variants. It is advised to use 2 µM of both oligo (dT) and random hexamers for cDNA synthesis when using a different kit.

qPCR reaction was set up with primers and probe sequences listed in Supplementary Table S4. Sso Advanced Universal Probes Supermix (Bio-Rad) was used following the manufacturer’s instructions. with the reactions were prepared in MicroAmp Endura 96-well plate and sealed with MicroAmp Optical Adhesive films, and the optical output was measured with StepOnePlus qPCR machine (all Applied Biosystems). A standard curve was included on the plate for each gene target tested; this standard curve was used to calculate cDNA copy number of the samples on the same plate. The FLuc or mScarlet-I cDNA copy number was used for normalization for the transformation efficiency. The resulting values from all the constructs were then normalized to the NLuc cDNA copy number from the *UBQ10:NLUC:NOS* in the same experiment, to account for any batch-to-batch variations.

### Functional dissection of the *HSP* and *FAD2* terminators

Five length deletion series were constructed for *HSP* and *FAD2* terminators. The deletions removed putative motifs that could affect their function to influence the gene expression level. Putative Stabilization/Destabilization Motifs^8^ while the rest were identified using RegRNA2.0^1^ and PASPA online tools^2^. Primers were designed to gradually remove functional sequence elements from the 3’ end of each terminator through PCR and subsequently cloned to mUAV. Site-directed mutagenesis by Gibson Assembly was employed to mutate the poly-A signal of the *HSP* Part3, while the same method was used to build *HSP* Part 5. Transformation of the construct was performed with PEG-transformation of Arabidopsis leaf mesophyll protoplasts in 96-well plates. The expression activity of the constructs was assayed using a plate reader, measuring the Nano luciferase activity normalized by Firefly luciferase. The construct we used was *UBQ10:Nluc:Terminator Part:UBQ10:Fluc:UBQ5*, housed in a pMAP vector.

### RNA folding prediction

For mRNA folding predictions, the transcription start site was predicted for each promoter by using TSSPlant^51^. The nucleotide sequence downstream of the TSS to the end of the terminator was used as the mRNA sequence. mRNA folding was predicted by using RNAFold software that uses Vienna RNA package.^50^ Thermodynamic ensemble prediction was used for folding energy predictions which accounts for all possible RNA secondary structures weighted by their Boltzmann probabilities, allowing the calculation of base pairing probabilities and ensemble free energy. The centroid algorithm was used for secondary structure prediction which is the structure with minimal base pair distance to all other secondary structures in the Boltzmann ensemble^50^.

## ACKNOWLEDGEMENTS

This project was funded by the Biotechnology and Biological Sciences Research Council (BBSRC) High-Value Compound from Plants (HVCfP) Network Proof-of-Concept Award (POC-NOV16-04), Royal Society University Research Fellowship (UF140640 and URF\R\201035), and Schmidt Science Polymath Award to NN, as well as the University of Edinburgh Principal’s Career Development PhD Studentship to AIA, the IBioIC CTP PhD Studentship to JN, and Imperial College Bioengineering Studentship to EGK.

